# Kctd15 regulates nephron segment development by repressing Tfap2a activity

**DOI:** 10.1101/2020.01.17.910760

**Authors:** Brooke E. Chambers, Eleanor G. Clark, Allison E. Gatz, Rebecca A. Wingert

## Abstract

A functional vertebrate kidney relies on structural units called nephrons, which are epithelial tubules that contain a sequence of segments each expressing a distinct repertoire of solute transporters. To date, the transcriptional codes driving regional specification, solute transporter program activation, and terminal differentiation of segment populations remain poorly understood. We demonstrate for the first time that the KCTD15 paralogs, *kctd15a* and *kctd15b*, function in concert to restrict distal early (DE)/thick ascending limb (TAL) segment lineage assignment in the developing zebrafish pronephros by repressing Tfap2a activity. During renal ontogeny, expression of these factors co-localized with *tfap2a* in distal tubule precursors. *kctd15* loss primed nephron cells to adopt distal fates by driving expansions in *slc12a1*, *kcnj1a.1*, and *stc1* marker expression. These phenotypes were resultant of Tfap2a hyperactivity, where *kctd15a/b*-deficient embryos exhibited increased abundance of this transcription factor. Interestingly, *tfap2a* reciprocally promoted *kctd15* transcription, unveiling a circuit of autoregulation operating in nephron progenitors. Concomitant *kctd15b* knockdown with *tfap2a* overexpression produced genetic synergy and further expanded the DE population. Our study provides strong evidence that a transcription factor-repressor feedback module employs tight regulation of Tfap2a and Kctd15 kinetics to control nephron segment fate choice and differentiation during kidney development.

## Introduction

Mammalian kidney organogenesis is unique in that it entails three waves of assembly, the pronephros, the mesonephros, and the metanephros, where each successive version becomes more morphologically complex than the previous.^1^ Other vertebrates, such as fish and amphibians, undergo two phases of kidney development, the final form being the mesonephros.^2^ Importantly, each kidney version across these animal kingdoms shares a conserved structural unit, the nephron. Recently, aquatic animal models such as fish and frogs have surfaced as powerful organisms to study the fundamentals of nephron development.^3^

The nephron is the functional unit of the kidney and is comprised of three core components: a glomerulus, an epithelial tubule, and a collecting duct. The tubule is compartmentalized into a series of proximal, intermediate, and distal segments. A key gap in knowledge remains in understanding the developmental signals necessary for the differentiation of segment-specific nephron cell types. More specifically, the genetic regulation of Loop of Henle (LOH) formation remains highly understudied. The LOH is partitioned into three limbs and initiates a concentration gradient by transporting water, sodium chloride, and potassium. Zebrafish, being an aquatic species, lack the first two limbs of the LOH because water conservation is not physiologically requisite. However, zebrafish do possess a distal early (DE) segment which is analogous to the mammalian thick ascending limb (TAL) of the LOH, which express a conserved suite of solute transporters including the orthologues of SLC12A1, KCNJ1, and CLCNK.^4, 5^ Defective TAL transporters are associated with neonatal and classical Bartter Syndrome, ion imbalance, polyuria, and renal failure. To this end, high-throughput screens in zebrafish have facilitated discovery of novel genes likely linked to renal tubular disorders.^6, 7^ More recently, a forward genetic screen and subsequent mutant analysis identified *transcription factor AP-2 alpha* (*tfap2a*) as a keystone regulator of the DE/TAL terminal differentiation program.^8^ TFAP2A belongs to the AP-2 family of transcription factors, which generally function in an embryonic context to control proliferation and differentiation.^9^

Two Tfap2a gene regulatory network (GRN) candidates are zebrafish paralogs *kctd15a* and *kctd15b*, which belong to the Potassium Channel Tetramerization Domain family. Proteins in this family harbor a conserved BTB/POZ (BR-C, ttk and bab/Pox virus and Zinc finger) protein-protein interaction motif situated at the N-terminus, however, demonstrate significant structural variability outside of this region. The BTB/POZ domain is essential for targeting proteins for ubiquitination and degradation.^10^ A diverse set of biological functions are assigned to these proteins such as transcriptional repression, gating of ion channels, regulating cytoskeletal elements, and acting as adaptor molecules. Mutations in KCTD genes can initiate human diseases such as breast cancer, medulloblastoma, epilepsy, pulmonary inflammation, and obesity highlighting the importance of further functional investigation.^11^ Humans, mice, and *Xenopus* possess one KCTD15 gene, however zebrafish acquired *kctd15a* and *kctd15b* paralogous versions due to an ancient species-specific genome duplication event. Zebrafish Kctd15a and Kctd15b amino acid residues exhibit a high degree of conservation with both splice variants of human KCTD15 protein and nearly identically alignment in the BTB/POZ functional motif (Figure S1).

In zebrafish, *kctd15a/b* inhibit neural crest development by two distinct modules: attenuation of canonical Wnt signaling and direct repression of Tfap2a. KCTD15 strongly inhibits TFAP2A by binding to its proline-rich activation domain.^12, 13^ Kctd15 is a substrate for SUMOylation; this post-translational modification is associated with transcriptional repression. However, previous studies suggest non-Sumoylated Kctd15 functions during neural crest formation.^14^ Genome wide association studies have revealed connections between Kctd15 and AP-2 to metabolic conditions such as obesity, diabetes, and eating disorders.^15, 16^ Contrary to neural crest development, in *Drosophila*, KCTD15 facilitates SUMOylation of Tfap2b to regulate consummatory behavior and repress the transduction of adipogenesis and insulin signaling pathways.^17, 18^ These findings authenticate the need for tissue-specific interrogation of Kctd15 function. Here, we demonstrate zebrafish *kctd15* paralogs are novel regulators of nephron segment commitment and inhibit DE/TAL differentiation by participating in repressor-mediated genetic feedback with *tfap2a*.

## Methods

### Ethics statement and zebrafish husbandry

Adult zebrafish were maintained at the University of Notre Dame Freimann Life Science Center. All studies were supervised by the University of Notre Dame Institutional Animal Care and Use Committee (IACUC), under protocol numbers 13-021 and 16-025. All WT experiments were conducted with the Tübingen strain. Embryonic zebrafish were incubated in E3 medium, staged and fixed as previously described.^19, 20^

### Whole-mount and fluorescent *in situ* hybridization (WISH, FISH)

WISH and FISH were performed as previously described.^21^ Digoxigenin and Fluorescein anti-sense RNA probes were synthesized by T7, T3, or SP6 *in vitro* transcription (Roche Diagnostics) from linearized IMAGE clone plasmids. Digoxigenin-labeled probes included: *kctd15a*, *kctd15b*, *kcnj1a.1*, *slc12a1*, *stc1, slc12a3,* and *trpm7*. Fluorescein-labeled probes included: *tfap2a*, *slc12a3*, *slc12a1*, *kcnj1a.1*, *cdh17*. Gene expression studies were performed in triplicate with sample size of n>20 for each replicate. Representative samples from each experimental group were imaged and analyzed.

### Whole-mount immunofluorescence (IF)

Whole mount IF was completed as previously described.^21^ For all IF experiments, embryos were fixed in freshly diluted 4% PFA for 2 hours at room temperature. 32% EM Grade PFA (Electron Microscopy Sciences, 15714) was diluted to 4% concentration in 1X PBS. After fixation, embryos were stored in 100% MeOH at −20°C for future use. Primary antibody dilutions consisted of anti-laminin (1:100; Sigma-Aldrich, L9393), anti-T4 supernatant (1:200; Developmental Studies Hybridoma Bank, Na-K-Cl cotransporter, AB_528406), and anti-Tfap2a (1:50; LifeSpan Biosciences, LS-C87212-100). Secondary antibodies (1:500) include goat anti-rabbit, rabbit anti-mouse, and donkey anti-goat (Invitrogen: A11034, A11061, A11055). 4,6-diamidino-2-phenylindole dihydrochloride (DAPI; Invitrogen, D1306) was used for nuclear staining.

### Morpholino knockdown and RT-PCR

Morpholino oligonucleotides (MO) were synthesized by GeneTools, LLC. MOs were solubilized in DNase/RNase-free water to make 4mM stock concentrations and stored at −20°C. Splice-blocking MOs for *kctd15a* and *kctd15b* genes were designed to target the exon1-intron1 splice sites, and microinjected at dosages of 1 ng and 3 ng, respectively. MO efficacy was validated by RT-PCR. RNA was extracted from pools of 20 embryos, cDNA was synthesized using random hexamers (Superscript IV, Invitrogen), and PCR was performed to amplify target site. Products were ran on a 1.5% Agarose gel, extracted, and sequenced. See Table S1 for specific MO and primer sequences.

### gRNA design and crispant generation

gRNAs targeting the BTB-POZ encoding regions in *kctd15a* and *kctd15b* were designed using the CHOPCHOP web-based tool (https://chopchop.cbu.uib.no/). *kctd15a* sgRNA1 and sgRNA2 targeted different regions in exon 3. *kctd15b* sgRNA1 and sgRNA2 targeted different regions in exon 1. gRNA templates were annealed to a constant oligonucleotide as previously described,^22^ and RNA was synthesized using the T7 Megascript kit (Ambion). Multiplexed gene editing was achieved by injecting a cocktail containing all four gRNAs. Microinjection mix was prepared by combining gRNAs (60 ng/μL) and Cas9 protein (0.8 μM) followed by incubation for 5 minutes at 37°C. Embryos were injected at the one-cell stage with ∼5 nL of injection mix. T7 endonuclease assay was used to confirm genome editing. For *kctd15a* and *kctd15b* crispant verification, primers were designed to flank both sgRNA target sites in exons 3 and 1 respectively. In short, DNA was prepared from individual animals and Accuprime Pfx SuperMix (Invitrogen) was used to amplify target sites. PCR products were column purified with QIAquick PCR purification kit (Qiagen). 300 ng of purified product and 2 μL 10x NEB Buffer 2 (total volume 20 μL) was rehybridized in a thermocycler using the following program: 5 min at 95°C, ramp down to 85°C at 2°C/s, ramp down to 25°C at 0.1°C/s, 25°C for 10 min. Rehybridized product was digested with 1 μL T7 endonuclease I enzyme (New England Biolabs) at 37°C for 1 hour and separated on a 1.5% agarose gel. See Table S1 for specific gRNA and primer sequences.

### Overexpression experiments

A *kctd15a* pCS2 construct was designed with Not1 and XhoI flanking the open reading frame allowing for *in vitro* synthesis of full-length sense cRNA using an SP6 mMessage Machine kit (Ambion). 60 pg of *kctd15a* cRNA was injected into one-cell stage embryos for overexpression. For *tfap2a* overexpression experiments, the *hs:tfap2a* transgenic [*Tg*(*hsp70:tfap2a*)^x24^], which was a generous gift from Bruce Riley (Texas A&M University, USA), was employed. To activate the heat-shock inducible *tfap2a* transgene, embryos were incubated at 38°C for 30 minutes at the 8ss or 12ss.^23, 24^ Genotyping for the *hs:tfap2a* allele was performed as previously described.^8^ See Table S1 for genotyping primers.

### Image Acquisition and statistical analysis

WISH samples were imaged with a Nikon Eclipse Ni with DS-Fi2 camera. FISH and IF samples were imaged with a Nikon C2 confocal microscope. The Nikon elements imaging software polyline tool was employed to quantify absolute lengths of gene expression domains. Confocal z-stacks were processed with Fiji (Image J) software and the plot profile feature was used to collect fluorescent intensity values. All graphical and statistical analyses were completed with GraphPad Prism. A minimum of three samples from control and experimental groups were imaged and scored. Unpaired t-test tests and one-way ANOVA analyses were conducted as appropriate and mean ± standard deviation was reported.

## Results

### *kctd15a* and *kctd15b* are expressed in the developing pronephros

Zebrafish and frog studies have previously reported *kctd15* transcript expression in the pronephros.^25, 26^ Data from the Genitourinary Development Molecular Anatomy Project (GUDMAP) detected KCTD15 expression in nascent mammalian nephrons. Microarray results indicated upregulated Kctd15 expression in developing renal vesicles in E12.5 mice (RID:Q-6PTG). Single cell RNA-sequencing of a week 17 human kidney cortex revealed elevated KCTD15 expression clustered with the developing nephron populace (RID:16-5HBT).^27, 28^ This documented expression positions *kctd15a* and *kctd15b* as excellent candidates of nephron regulation. To expand on these previous studies, we assembled a comprehensive time course of *kctd15a* and *kctd15b* spatial expression over the span of kidney development using the genetically tractable zebrafish pronephros model. Upon WISH analysis of *kctd15a* and *kctd15b* in wild-type (WT) animals, transcripts were first detectable in the intermediate mesoderm (IM) at the 10 somite stage (ss) (Figure 1A). Throughout the duration of pronephric development, *kctd15a* and *kctd15b* showed similar expression patterns becoming increasingly restricted to the distal nephron progenitor domain (Figure 1B). Because the expression patterns of these factors in the developing kidney show minimal deviation from one another, it is highly possible they function redundantly in this context.

**Fig. 1.**
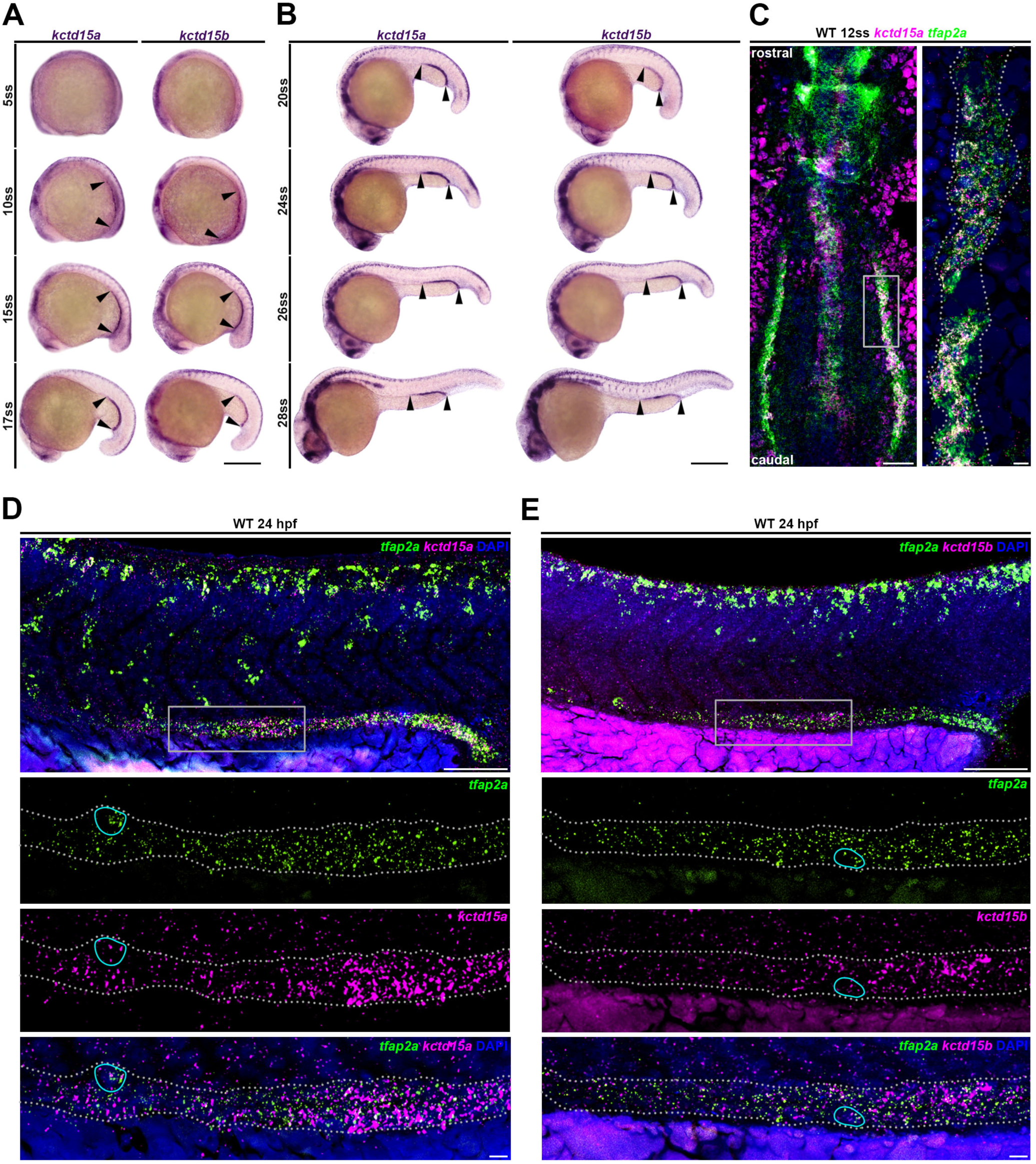
*kctd15a* and *kctd15b* are expressed in developing distal nephron precursors. **(A)** WISH of *kctd15a* and *kctd15b* during WT embryogenesis. Black arrowheads mark expression within developing renal field. Scale bar: 200 µm. **(B)** WISH of *kctd15a* and *kctd15b*. Black arrowheads mark expression within distal nephron precursors. Scale bar: 200 µm. **(C)** FISH of *kctd15a* (magenta) and *tfap2a* (green) in 12 ss WT flatmount. Grey box indicates featured region in right panel. Grey dots demarcate renal progenitor stripe. Scale bars: 100 µm (left), 5 µm (right). **(D)** FISH of *tfap2a* (green) and *kctd15a* (magenta) at 24 hpf. Grey box represents region imaged at higher magnification (bottom panels). Cyan encircles an example of a co-expressing cell. Scale bars: 35 µm (top), 5 µm (bottom). **(E)** FISH of *tfap2a* (green) and *kctd15b* (magenta) at 24 hpf. Grey box represents region imaged at higher magnification (bottom panels). Cyan encircles an example of a co-expressing cell. Scale bars: 35 µm (top), 5 µm (bottom).

Next, we wanted to determine how the *kctd15a* and *kctd15b* expression domains spatially correlate to *tfap2a* transcripts in the developing pronephros, since these factors repress Tfap2a in neural crest.^12, 25^ Using fluorescent *in situ* hybridization (FISH), *kctd15a* transcripts were found to be co-expressed throughout the *tfap2a^+^* IM populace at the 12 ss in WT embryos. (Figure 1C). At 24 hours post fertilization (hpf), *kctd15a* and *kctd15b* were expressed throughout the *tfap2a^+^* pronephric domain (Figure 1D,E). These experiments demonstrate *kctd15a* and *kctd15b* are expressed in developing distal precursors and their localization largely overlaps with *tfap2a* at multiple timepoints during development.

### *kctd15a/b* regulates the expression of distal nephron markers

In zebrafish, *kctd15a/b* deficiency leads to an expansion of neural crest fate,^25, 29^ most likely due to the inability to repress Tfap2a. Because *kctd15a/b* function had never been studied in the context of kidney organogenesis, we developed loss and gain of function strategies to assess their involvement in nephrogenesis. Knockdown of *kctd15a* and *kctd15b* was achieved by injecting morpholinos to disrupt splicing between the exon 1 and exon 2 junctions, and validated by RT-PCR analyses (Figure S2). For a parallel independent loss of function model, we genetically engineered F0 *kctd15a, kctd15b, and kctd15a/b* CRISPR mutants (crispants) by multiplexing sgRNAs targeting different regions of the BTB/POZ functional domain to induce biallelic disruptions (Figure S3). T7 endonuclease assays and Sanger sequencing confirmed successful genome editing (Figure S3). *kctd15a/b* morphants, *kctd15a* crispants, and *kctd15b* crispants developed pericardial edema and increased dorsal head pigmentation by 48 hpf consistent with other live phenotype reports (Figure S2, S3).^25, 29^

Because *kctd15* transcripts localized to the developing pandistal pronephric region (Figure 1), we first decided to survey for alterations in DE and DL marker expression, which are analogous to the mammalian TAL and distal convoluted tubule (DCT), respectively. *kctd15a*, *kctd15b*, and *kctd15a/b* morphants and crispants exhibited drastic expansions of the DE markers *kcnj1a.1* and *slc12a1* as compared to WT controls (Figure 2A-D). In each *kctd15*-deficient state, *slc12a1^+^* cells encroached the DL territory marked by *slc12a3* (Figure 2C). Notably, previous study indicated when *tfap2a* is overexpressed, an expansion of DE solute transporter expression occurs.^8^ These parallel phenotypes highlight the possibility that this alteration of distal nephron fate could be due to inadequate repression of Tfap2a protein by Kctd15a/b. Generally, *kctd15* crispants manifested slightly milder DE phenotypes as compared to *kctd15* morphants, most likely because F0 crispant cellular composition is mosaic in nature and generates a mix of homozygous and heterozygous mutants. Nevertheless, *kctd15* crispants recapitulated *kctd15* morphant distal nephron phenotypes corroborating the use of MO mediated knockdown in subsequent experiments. As expected, compound MO knockdown of *kctd15a* and *kctd15b* elicited the most severe expansions of DE marker expression, suggesting that these paralogs evolved redundant functions in teleosts (Figure 2B, 2D). Overexpression of *kctd15a* cRNA produced a significantly decreased DE in conjunction with an expanded DL segment (Figure 2A-D). *kctd15a* overexpression also caused a substantial decline in *kcnj1a.1*^+^ ionocyte number (Figure S4). Matching phenotypes occurred upon *kctd15b* overexpression (data not shown). Next, we assessed *Slc12a1* protein expression in *kctd15a/b* morphants. We found that *kctd15a/b* morphants exhibit a significant expansion of the *Slc12a1* protein domain compared to WT embryos (Figure 2E). Generation of fluorescent intensity profiles confirmed a significant elevation in *Slc12a1* signal in *kctd15a/b*-deficient embryos compared to WT controls (Figure 2E-H). Quantification of cell number found that *kctd15a/b* morphants have nearly double the number of *Slc12a1*^+^ cells (Figure 2I).

**Fig. 2.**
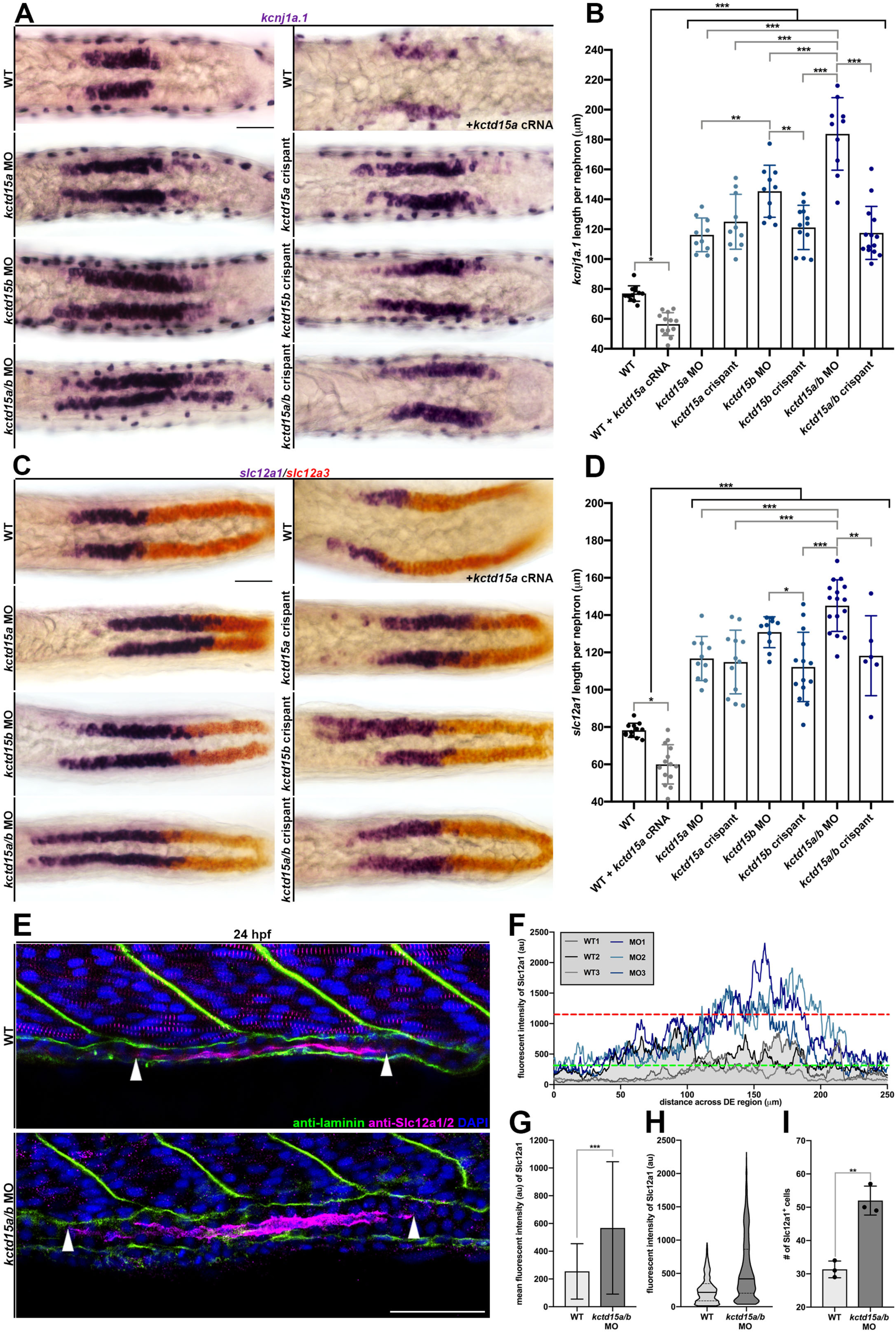
*kctd15a/b* loss-of-function initiates expansion of DE lineage markers. **(A)** WISH of *kcnj1a.1* (purple) at 24 hpf in WT, WT + *kctd15a* cRNA, *kctd15* MO, and *kctd15* crispants. Scale bar: 35 µm. **(B)** Absolute length quantification of *kcnj1a.1*. Black bracket clusters groups together for collective comparison. **(C)** WISH of *slc12a1* (purple) and *slc12a3* (red) at 24 hpf in WT, WT + *kctd15a* cRNA, *kctd15* MO, and *kctd15* crispants. Scale bar: 35 µm. **(D)** Absolute length quantification of *slc12a1*. Black bracket clusters groups together for collective comparison. **(E)** IF of Slc12a1/2 (magenta) and laminin (green) in WT and *kctd15a/b* MO at 24 hpf. White arrowheads mark limits of Slc12a1/2 pronephric expression. Scale bar: 35 µm. **(F)** Fluorescent intensity plot of 3 WT (greyscale) and 3 *kctd15a/b* MO individuals (blue). Green dashed line equates to WT Slc12a1/2 intensity threshold (au). Red dashed line represents WT maximum Slc12a1/2 intensity value (au). **(G)** Mean Slc12a1/2 fluorescent intensity graph. **(H)** Violin plot displaying point distribution of Slc12a1/2 intensity values collected. **(I)** Quantification of number of Slc12a1^+^ pronephric cells. n≥3. *P<0.05; **P<0.01; ***P<0.001. Data are mean ± s.d. Absolute lengths compared by ANOVA. Mean fluorescent intensity and cell counts analyzed by unpaired t-tests.

Further, we examined a distal nephron associated cell type, the corpuscle of Stannius (CS). Briefly, the CS is an aggregate of cells situated between the DE and DL segments that eventually bud off the pronephric tubule to form an endocrine gland that maintains calcium homeostasis.^30^ A recent study illustrated that CS cells transdifferentiate from the DE segment by a gland extrusion mechanism.^31^ *kctd15a*, *kctd15b*, and *kctd15a/b* morphants and crispants all possess elevated numbers of CS cells as marked by *stc1* compared to WTs (Figure 3A,B). Surprisingly, we noticed ectopic *stc1^+^* cells formed at the proximal end of the DE domain that were clearly separate from the principal CS cell cluster in *kctd15a/b* morphant embryos (Figure 3A). Upon closer examination, we discovered *kctd15a/b* morphants undergo premature CS gland extrusion at 24 hpf, as this group of cells formed a basement membrane and separated from the pronephric tubule (Figure 3C). As *tfap2a* deficiency affects CS differentiation, it is not surprising that disrupting the balance of *kctd15a/b* alters this process as well.^8^ Collectively, our genetic experiments revealed alterations in DE, DL, and CS cell differentiation, identifying *kctd15* as a novel regulator of distal nephron identity.

**Fig. 3.**
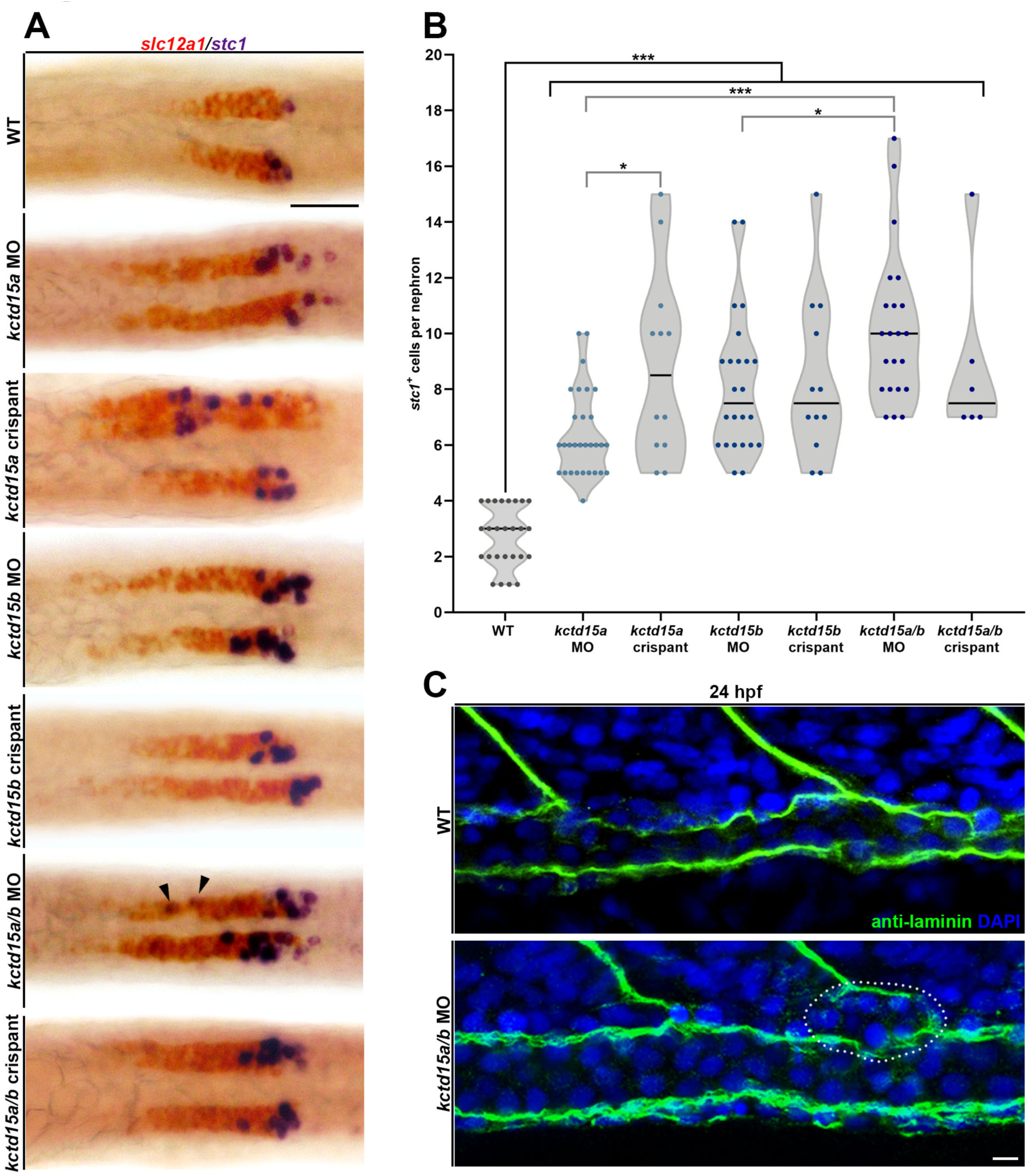
*kctd15a/b*-deficiency elevates CS differentiation. **(A)** WISH of *slc12a1* (red) and *stc1* (purple) at 24 hpf in WT, *kctd15* MO, and *kctd15* crispants. Black arrowheads indicate proximal stray *stc1*+ cells clearly separate from the main CS cluster in *kctd15a/b* MO. Scale bar: 35 µm. **(B)** Quantification of *stc1*^+^ cell number per nephron. Black bracket clusters groups together for collective comparison. **(C)** IF of laminin (green) in 24 hpf WT and *kctd15a/b* MO. Premature basement membrane formation occurs in *kctd15a/b* morphant, which separates the budding CS cell cluster (white dotted circle) from the pronephric tubule. Scale bar: 5 µm. n≥3. *P<0.05; **P<0.01; ***P<0.001. Data are mean ± s.d. Cell counts compared by ANOVA.

### *kctd15a/b* depletion initiates ectopic DE formation in flanking pronephros segments

Next, we decided to examine the effect of dual *kctd15a/b* knockdown on the neighboring nephron segment populations, as zebrafish *kctd15a* and *kctd15b* share over 90 percent amino acid sequence identity and likely have overlapping functions (Figure S1). We employed FISH to visualize the DE and DL nephron segment domains marked by *kcnj1a.1* and *slc12a3* respectively. In 24 hpf WT animals, there is a sharp separation between the DE and DL segment compartments (Figure 4A,B). Conversely, in *kctd15a/b* morphants, *kcnj1a.1*^+^ cells were located in the DL segment. This significant overlap of DE and DL segment identities was made evident upon generation of a fluorescent intensity plot depicting a representative *kctd15a/b* morphant (Figure 4C). A portion of these cells dually expressed *kcnj1a.1* and *slc12a3*, suggesting segment fate infidelity (Figure 4A). This dual marker expression is highly reminiscent of the resultant phenotype triggered by overexpression of *tfap2a*.^8^ Absolute length quantification of *slc12a3* revealed that the *kctd15a/b* morphant DL domain is significantly shorter than WT controls (Figure 4D). These data indicate that *kctd15a/b* deficiency results in an expansion of the DE lineage at the expense of the DL. To probe the rostral DE boundary and its adjacent proximal straight tubule (PST) segment, *trpm7* expression was assessed. WT animals have a well-defined DE/PST boundary (Figure 4E,F). Contrary to this, *kctd15a/b* morphant DE cells occupied the *trpm7^+^* PST region (Figure 4E,G). Additionally, some cells co-expressed *kcnj1a.1* and *trpm7*, indicating *kctd15a/b*-deficiency was potent enough to induce proximal nephron cells to express a distal nephron signature gene. Further, the absolute length of the PST was significantly reduced in *kctd15a/b* morphants (Figure 4H). Taken together, these data suggest that *kctd15a/b* factors function to suppress DE lineage in neighboring segment populations during nephrogenesis.

**Fig. 4.**
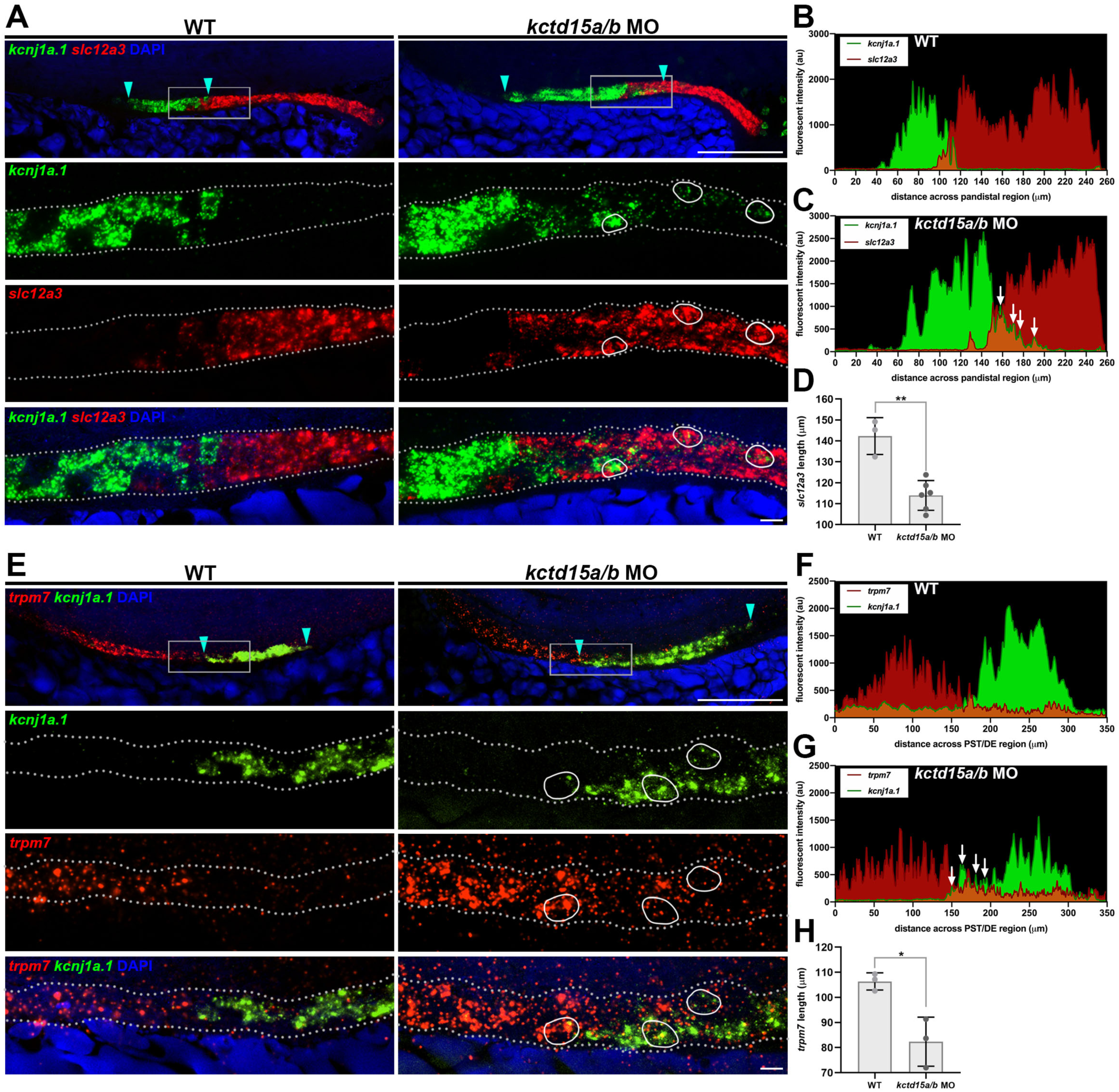
Loss of *kctd15a/b* sways neighboring pronephric fates to express DE signature. **(A)** FISH of *kcnj1a.1* (green) and *slc12a3* (red) at 24 hpf in WT and *kctd15a/b* MO. Cyan arrowheads denote *kcnj1a.1* boundaries. Grey box indicates area featured at higher magnification in panel below. White circles represent individual cells dually expressing *kcnj1a.1* and *slc12a3*. Scale bars: 70 µm (top), 5 µm (bottom). **(B)** WT fluorescent intensity plot of *kcnj1a.1* (green) and *slc12a3* (red). **(C)** *kctd15a/b* MO fluorescent intensity plot of *kcnj1a.1* (green) slc12a3 (red). White arrows indicate ectopic *kcnj1a.1* signal invading the *slc12a3* domain. **(D)** Absolute *slc12a3* length quantification. **(E)** FISH of *trpm7* (red) and *kcnj1a.1* (green) at 24 hpf in WT and *kctd15a/b* MO. Cyan arrowheads denote *kcnj1a.1* boundaries. Grey box indicates area featured at higher magnification in below panel. White circles represent individual cells dually expressing *kcnj1a.1* and *trpm7*. Scale bars: 70 µm (top), 5 µm (bottom). **(F)** WT fluorescent intensity plot of *trpm7* (red) and *kcnj1a.1* (green). **(G)** *kctd15a/b* MO fluorescent intensity plot of *trpm7* (red) and *kcnj1a.1* (green). White arrows indicate ectopic *kcnj1a.1* signal invading the *trpm7* domain. **(H)** Absolute *trpm7* length quantification. n≥3. *P<0.05; **P<0.01. Data are mean ± s.d. Absolute lengths compared by unpaired t-tests.

### *kctd15a/b* knockdown expands Tfap2a expression in the developing kidney

To evaluate if *kctd15a/b* are regulating nephron differentiation by affecting Tfap2a localization or stability, we performed IF experiments on 24 hpf *kctd15a/b* morphants. *kctd15a/b* morphants showed expanded pronephric expression of Tfap2a protein compared to WTs (Figure 5A). Specifically, this morphant expansion was characterized by residual proximal nephron Tfap2a protein expression. This greatly contrasts WTs that exhibit an abrupt cutoff of Tfap2a protein expression at the proximal limit of the Slc12a1^+^ DE (Figure 5A). Generation of fluorescent intensity profiles revealed differentially elevated morphant Tfap2a expression that was particularly evident within a 100-220 µm region that maps to the DE territory (Figure 5B). *kctd15a/b* morphants also exhibited a significant elevation of Tfap2a fluorescent signal within the 100-220 µm zone (Figure 5C,D). Further, *kctd15a/b* morphants exhibited a statistically significant increase in Tfap2a^+^ pronephric nuclei than WT controls (Figure 5E). In summary, these results indicate that when *kctd15a/b* repressors are absent, the number of Tfap2a expressing cells increases in the pronephros and simultaneously parallels an expansion of the Slc12a1^+^ DE. Our data illustrating alterations in pronephric Tfap2a protein expression reveals a new mechanistic layer that is distinct from previous *in vitro* studies, which found nuclear extracts exhibited no changes in Tfap2a protein abundance in the absence of Kctd15 and the main mode of inhibition was executed by direct binding to the Tfap2a transactivation domain.^12^ Whole-mount IF allowed for tracking cellular changes in Tfap2a protein expression in specific nephron regions. Based on these IF results, we hypothesized that expanded Tfap2a pronephric expression is the root cause of the DE lineage alterations that occur in *kctd15a/b* morphants.

**Fig. 5.**
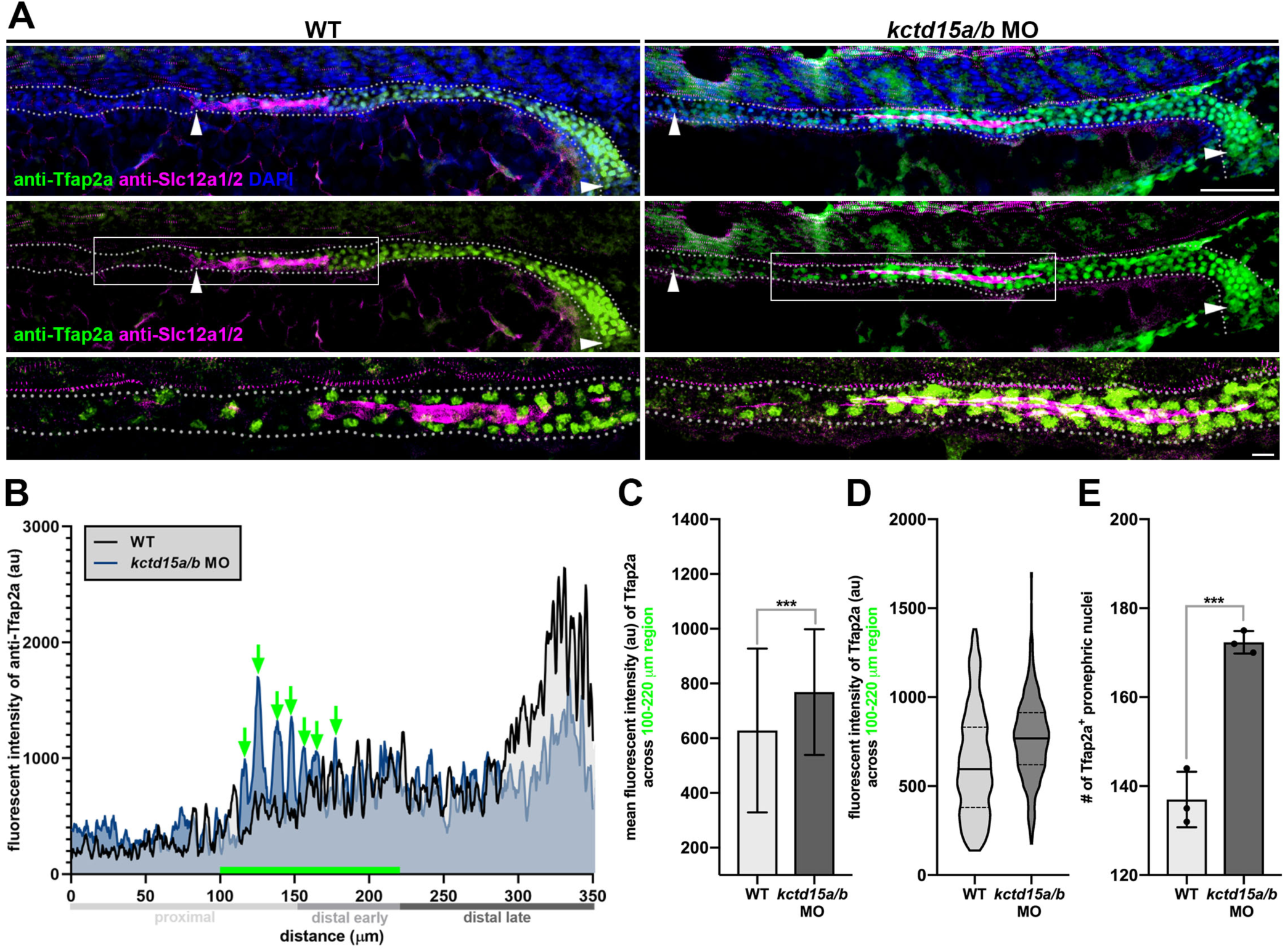
*kctd15a/b* knockdown expands Tfap2a protein expression in the pronephros. **(A)** Whole-mount IF of Tfap2a and Slc12a1/2 in WT and *kctd15a/b* MO. Scale bars: 35 µm (top), 5 µm (bottom). White arrowheads indicate limits of pronephric Tfap2a expression domain. Grey box highlights region at higher magnification below. Grey dotted lines demarcate pronephric tubule. **(B)** Fluorescent intensity plot of Tfap2a featuring one representative WT and *kctd15a/b* MO sample. Green arrows label differential Tfap2a signal peaks. Green bar spans region of differential expression corresponding to DE locale (100-220 µm). **(C)** Mean fluorescent intensity (au) graph of Tfap2a across the 100-220 µm region. **(D)** Violin plot displaying point distribution of Tfap2a intensity values collected across the 100-220 µm region. **(E)** Quantification of number of Tfap2a^+^ pronephric cells. n≥3. ***P<0.001. Data are mean ± s.d. Mean fluorescent intensity and cell counts analyzed by unpaired t-tests.

### *kctd15a/b* and *tfap2a* engage in genetic feedback circuitry

Because we discovered heightened Tfap2a protein signal in the pronephros in response to *kctd15a/b* deficiency (Figure 5), we postulated a genetic feedback mechanism might be responsible for this phenotype. In support of this concept, a previous study reported *kctd15a/b* zebrafish mutants manifest upregulated *tfap2a* neural expression at the 8-9 ss.^29^ To probe potential genetic feedback processes, we performed FISH at the 10 ss to detect fluctuations in *tfap2a* transcript expression. *kctd15a/b* knockdown led to measurable expansions of *tfap2a* mRNA expression within the IM populace, which is comprised of renal progenitors that give rise to the pronephros (Figure 6A). In WT flatmounts, the rostral end of *tfap2a* IM expression aligns with the somite 8 landmark. However, in *kctd15a/b* morphants, this expression domain extends in the rostral direction well beyond somite 8. We also noted elevated *tfap2a* expression in *kctd15a/b*-deficient hindbrain regions (Figure 6A). Then, we surveyed *tfap2a* IM fluorescent intensity and standardized the region of data collection by tracking values from somite 7 to the end of the pre-somitic mesoderm. Plot overlay of *tfap2a* fluorescent intensity profiles from representative samples revealed morphant IM signal was consistently elevated above WT levels (Figure 6B). Further, *tfap2a* mean fluorescent intensity was significantly higher in morphants compared to WT controls (Figure 6C,D). This *tfap2a* transcript pulse equilibrates in the pronephros as embryonic development progresses, as normal levels of mRNA were detected in *kctd15a/b* morphants and crispants by 24 hpf (data not shown).

**Fig. 6.**
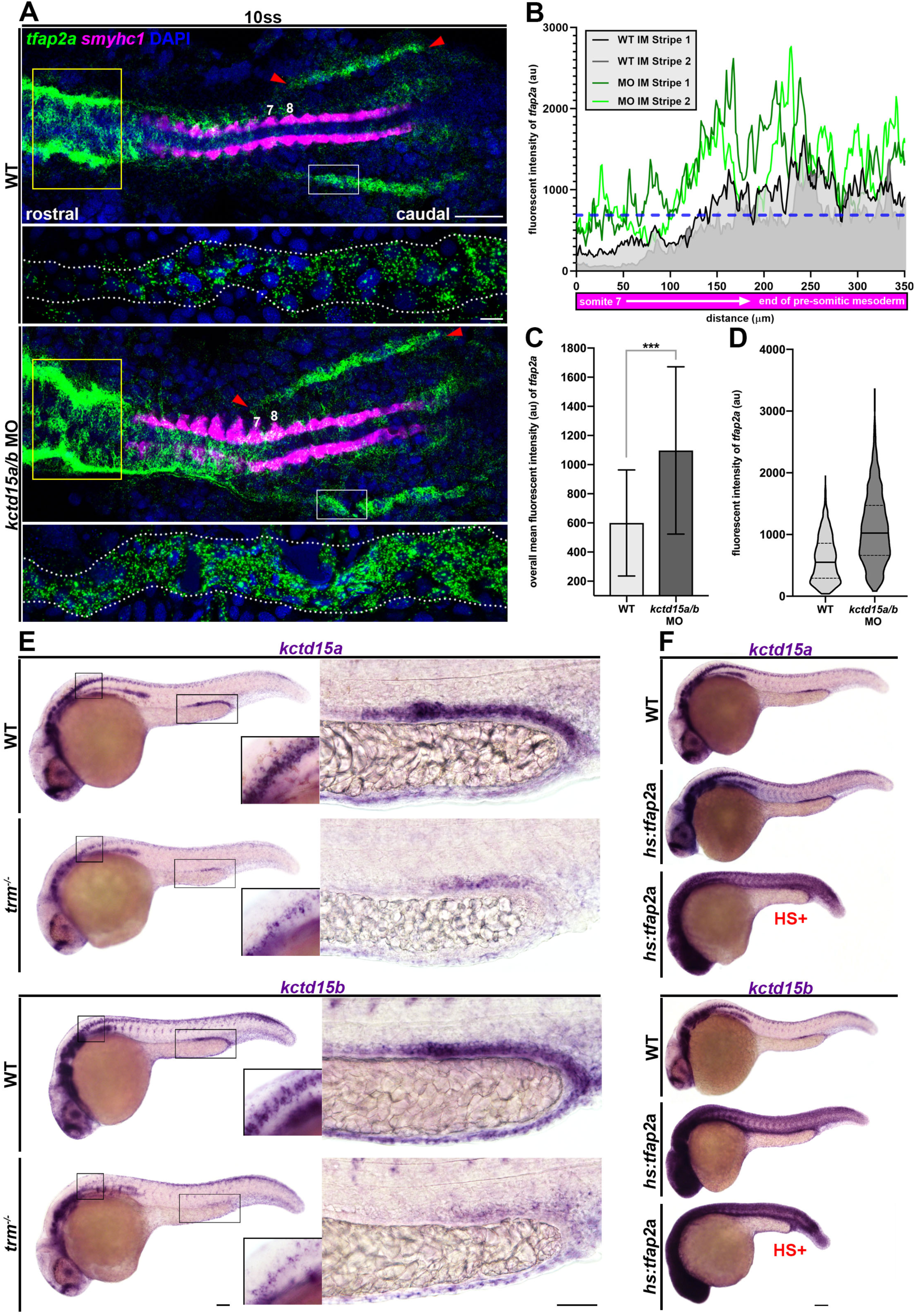
*kctd15a/b* and *tfap2a* participate in genetic crosstalk. **(A)** Whole-mount FISH of *tfap2a* (green) and *smyhc1* (magenta) in 10 ss WT and *kctd15a/b* MO flatmounts. Yellow box indicates differential *tfap2a* hindbrain expression. Red arrows demarcate *tfap2a* IM stripe expression limits. Numbers label somite 7 and 8 landmarks. White box indicates region depicted in below panel at higher magnification. Scale bars = 100 µm (above), 5 µm (below). **(B)** Fluorescent intensity plot of *tfap2a* IM expression featuring one representative WT (greyscale) and *kctd15a/b* MO (green) sample. Blue dotted line signifies WT mean fluorescent intensity threshold. **(C)** Overall mean fluorescent intensity (au) graph of *tfap2a* IM expression. Fluorescent intensity values were collected from somite 7 to the end of the pre-somitic mesoderm. **(D)** Violin plot displaying point distribution of *tfap2a* intensity values. **(E)** WISH of *kctd15a* or *kctd15b* (purple) in WT and *trm^-/-^* mutants. Inset features neural expression. Right panel features pronephric expression. Scale bars: 70 µm (left), 35 µm (right). **(F)** WISH of *kctd15a* or *kctd15b* (purple) in WT and *hs:tfap2a*. HS+ (red) signifies heat-shock treatment at the 8ss. Scale bar: 70 µm. n≥3. ***P<0.001. Data are mean ± s.d. Mean fluorescent intensity analyzed by unpaired t-tests.

Next, we sought to determine if *kctd15a* and *kctd15b* expression are affected by *tfap2a*-deficiency. *trm* mutants exhibited considerable reductions in *kctd15a* and *kctd15b* mRNA expression at 24 hpf as compared to WT controls (Figure 6E). These visual reductions in *kctd15a/b* expression were observed in both neural and pronephric tissues. Then we tested if overexpression of *tfap2a* via a heat-shock inducible transgene is sufficient to activate *kctd15a/b* transcription. *hs:tfap2a* animals that underwent heat-shock treatment at the 8ss displayed a global escalation of *kctd15a* and *kctd15b* expression (Figure 6F). Surprisingly, *hs:tfap2a* control animals that did not undergo heat-shock treatment also exhibited increased overall *kctd15a* and *kctd15b* expression, but to a lesser extent than heat-shock treated siblings (Figure 6F). We believe this is an artifact of a leaky *tfap2a* transgene. Nonetheless, this data illustrates that *kctd15a/b* transcript expression is extremely sensitive to *tfap2a* dosage. Altogether, our findings expose a previously undescribed level of genetic interaction where *tfap2a* regulates the transcription of its own repressors *kctd15a* and *kctd15b*. Excitingly, these results divulged an autoregulatory feedback loop operating in the context of nephron differentiation where *tfap2a* controls its own abundance through the *kctd15a*/*b* genetic circuit.

### *kctd15a/b* counterbalances *tfap2a* to restrict DE differentiation during pronephros segmentation

To further substantiate that *kctd15a/b* functions in nephron development by repressing Tfap2a activity, we disrupted the stoichiometry of these factors by concomitant *kctd15b* knockdown and induced *tfap2a* overexpression at the 12 ss via the heat-shock transgene. Interestingly, *tfap2a* overexpression produced significantly shorter *cdh17*^+^ tubules (Figure A,B). To account for these truncated tubules in subsequent analyses, we calculated a ratio, DE length:tubule length, to statistically compare all groups. We found no significant difference in DE:tubule percentage between WT and *hs:tfap2a^+/-^* controls (Figure 7A,C). The DE fraction was similarly increased in WT + *kctd15b* MO, *hs:tfap2a^+/-^* + *kctd15b* MO, and *hs:tfap2a* HS+ groups. Remarkably, pairing *kctd15b* knockdown with *tfap2a* overexpression initiated a drastic expansion of DE marker expression that was more severe than all other experimental groups. In these dual-treated animals, ectopic *slc12a1*^+^ cells formed in both proximal and distal nephron segments (Figure 7A). These synergistic phenotypes upon concomitant knockdown of *kctd15b* and overexpression of *tfap2a* corroborate the interdependence of these factors during nephron segmentation. In sum, our genetic studies divulge *kctd15* paralogs as essential repressors that function during nephrogenesis to stabilize the DE lineage by antagonizing the *tfap2a* differentiation pathway.

**Fig. 7.**
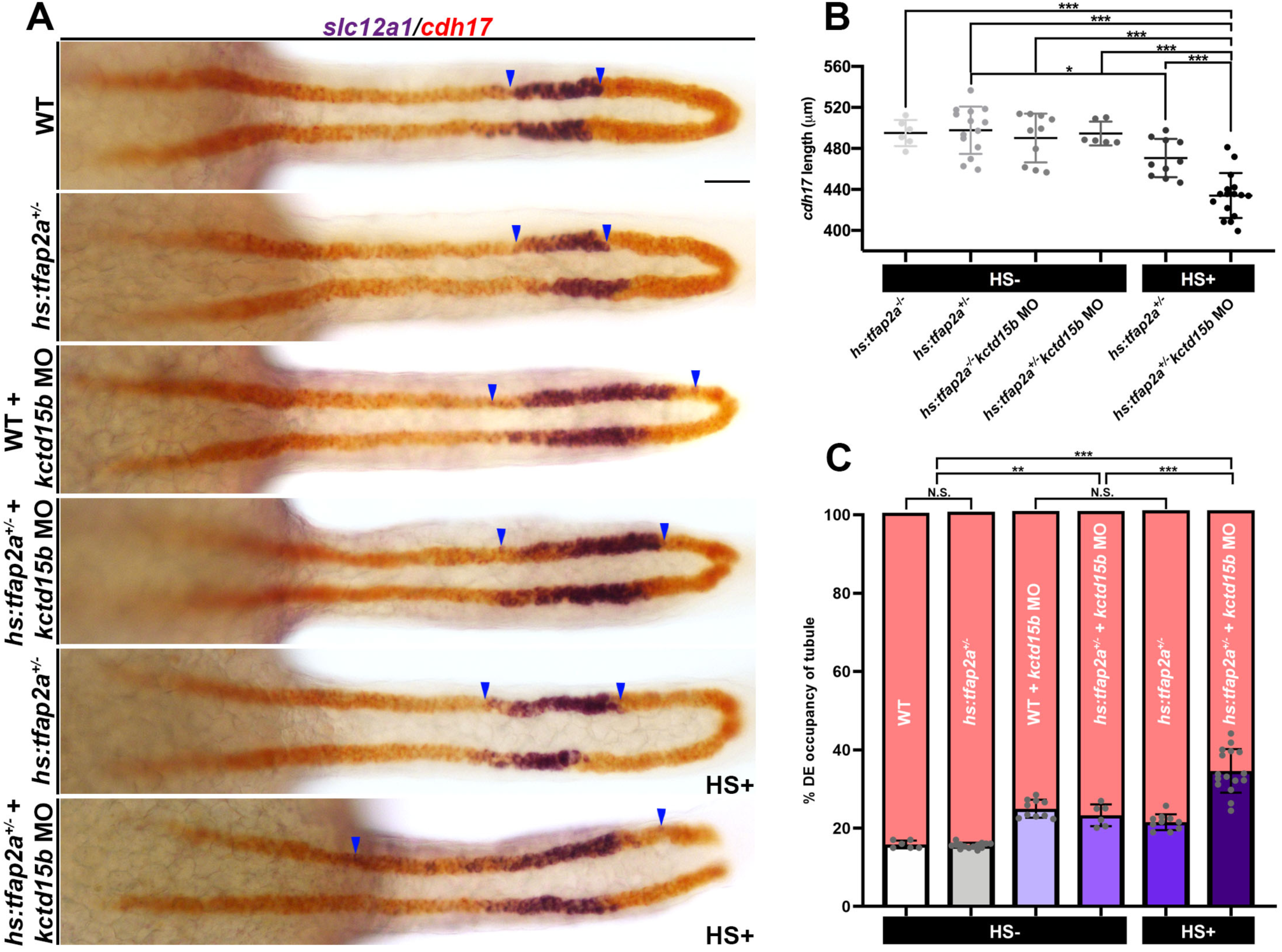
*kctd15a/b*-*tfap2a* autoregulatory feedback loop balances DE pronephric differentiation. **(A)** WISH of *slc12a1* (DE, purple) and *cdh17* (tubule, red) in *hs:tfap2a* transgenic background in combination with *kctd15b* MO treatment. HS+ indicates application of heat-shock treatment at the 12ss. Blue arrowheads annotate proximal and distal edges of *slc12a1* expression. Scale bar: 35 µm. **(B)** Quantification of *cdh17* length in control and treatment groups. **(C)** Bar graph depicting percent DE occupancy of the tubule. HS-= no heat-shock treatment, HS+ = heat-shock treatment. n≥6. *P<0.05; ***P<0.001. Data are mean ± s.d. Absolute lengths and percentages compared by ANOVA.

## Discussion

During embryonic development, Tfap2 genes are dose-dependent transcription factors, therefore require tight regulation.^8, 32–35^ Here, our data supports a new model where *kctd15* fine-tunes Tfap2a activity to control nephron segmentation. We found *kctd15a/b-*deficient nephrons exhibited expanded DE and CS lineages (Figure 8). This surge in DE and CS fate occurred at the expense of neighboring segments, as PST and DL marker expression were concomitantly decreased. *kctd15* loss initiated ectopic expression of DE signature genes in adjacent nephron compartments. Frequently, we detected a fraction of cells exhibiting dual segment marker expression insinuating partial fate conversion. This phenomenon of mixed segment profiles was previously documented in Abd-B Hox-deficient and *tfap2a* overexpressing nephrons.^8, 36^ Previous *in vitro* experiments established a direct mode of interaction where KCTD15 inhibits AP-2 in whole zebrafish embryo and HEK293T lysates.^12^ In light of these mammalian cell line and zebrafish studies, we suspected *kctd15*-deficient nephron phenotypes were caused by increased Tfap2a activity. Interestingly, elevated Tfap2a protein abundance was detected in *kctd15*-deficient pronephroi, which diverges from previous neural crest studies. We believe accumulation of Tfap2a protein is a consequence of genetic feedback circuitry because 1) *kctd15* loss expanded *tfap2a* mRNA expression in developing renal progenitors and 2) *tfap2a* promotes *kctd15a* and *kctd15b* transcription. In accordance with these observations, Tfap2a can promote its own transcription by binding to an autoregulatory element.^37^ If Tfap2a is unable to be repressed, positive autoregulation could explain heightened *tfap2a* transcript production in response to *kctd15* loss-of-function.

**Fig. 8.**
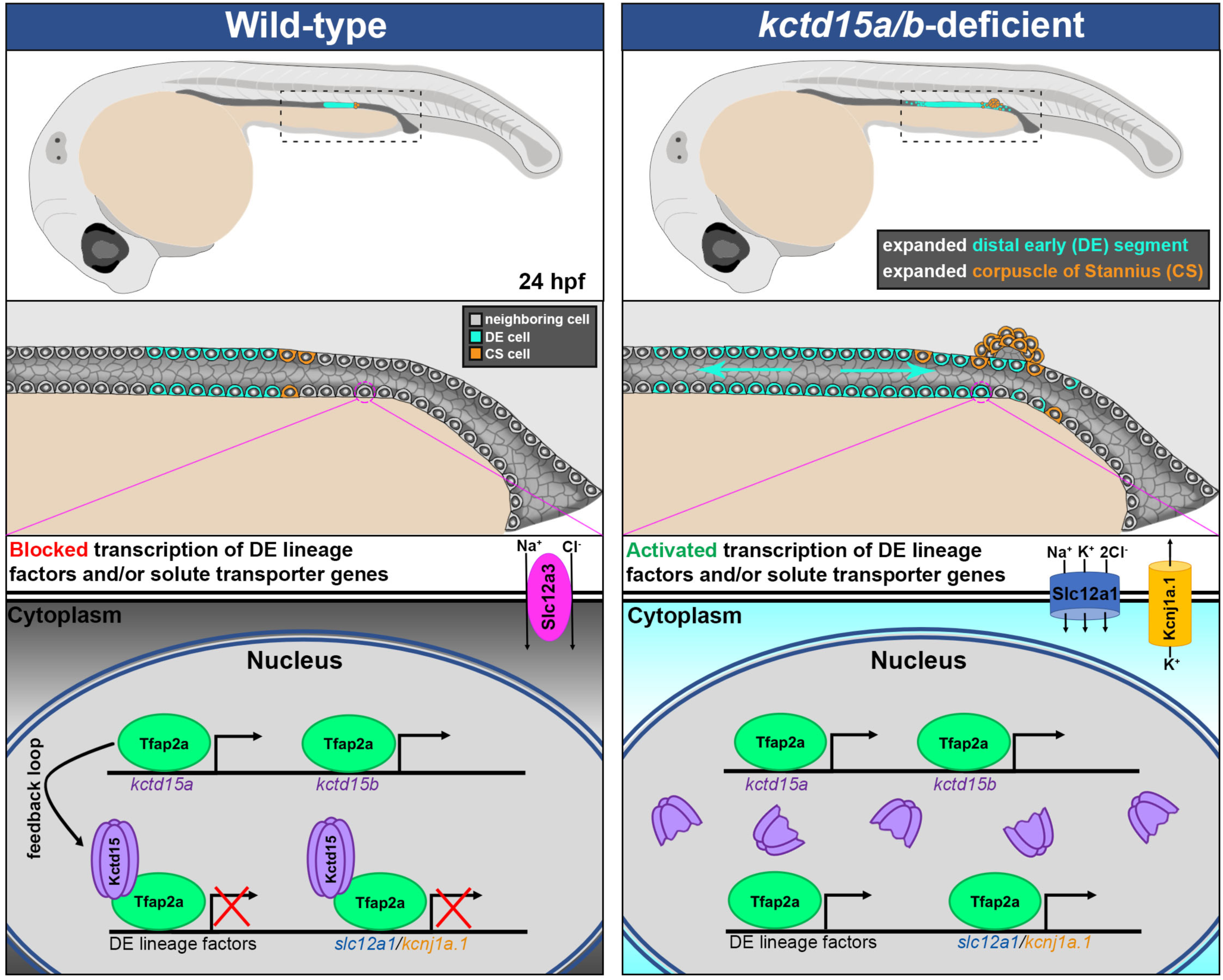
Working model illustrating the proposed role of the Kctd15-Tfap2a interaction in maturing nephron cells. WT (left), *kctd15a/b*-deficient context (right) at 24 hpf. Distal early (DE, cyan), corpuscle of Stannius (CS, orange), neighboring cell populations (grey). Black dashed box indicates featured distal nephron zone. Cyan arrows signify expansion of DE differentiation in *kctd15a/b*-deficient nephron. Magenta circle outlines single cell featured in bottom panel. In WT, Tfap2a promotes *kctd15* expression at the transcriptional level and Kctd15 protein (purple) represses Tfap2a (green) activity, allowing for proper nephron segment differentiation. In *kctd15*-deficiency, predicted truncated Kctd15 protein is not able to inhibit Tfap2a activity in neighboring cell populations, allowing for ectopic activation of DE lineage factors and solute transporters (*slc12a1*, *kcnj1a.1*).

Alternatively, it is feasible excess pronephric Tfap2a protein is present because it is not properly targeted for degradation. Previous studies have demonstrated KCTD proteins can function as adaptor molecules for E3 ubiquitin ligases to facilitate post-translational modifications of bound protein interactors.^38^ Further, KCTD family members are substrates for SUMOylation and ubiquitinylation, which can alter protein stability and initiate degradation respectively. There is evidence for SUMO-conjugation of TFAP2A, which suppresses transcriptional activation and prevents mesenchymal-to-epithelial transition in breast cancer cells.^39–42^ Further investigation is needed to determine whether Kctd15 facilitates SUMOylation of Tfap2a in nephron precursors. Nonetheless, our *kctd15a/b*-deficient genetic manipulations truncate the BTB/POZ domain and would prevent Kctd15-Tfap2a direct interactions from occurring. Because feedback circuitry is disrupted, Tfap2a activity is left unchecked in neighboring nephron segments, allowing downstream targets like DE-specific lineage factors and solute transporters to be ectopically expressed (Figure 8). In support of our model, *kctd15b* knockdown coupled with *tfap2a* overexpression yielded a synergistic phenotype where nephron cell composition was shifted toward DE assignment. In conclusion, our studies reveal a novel feedback circuit operating within the context of kidney development, where a *tfap2a-kctd15* transcription factor-repressor module programs DE/TAL differentiation.

Our studies help to fill a key void in the literature by addressing developmental signaling required for DE/TAL formation. Due to the anatomical constraints of studying mammalian nephron development *in vivo*, there is a limited knowledge of cell diversity and dynamics within the maturing LOH anlagen. Zebrafish studies provide a complementary launch point to explore the how functionality of candidate factors, such as *tfap2a* and *kctd15*, create nephron cell-types in vertebrates. Recently, scRNA-seq and fate-mapping in mice revealed early forming LOH (juxtamedullary) and late forming LOH (cortical) have distinct gene signatures.^43^ LOH outgrowth from the S-shaped tubule is adaptive and oriented by long-range cues from medullary collecting ducts, however signaling pathways navigating this process have yet to be identified.^44^ Current efforts to advance personalized medicine by building patient-derived kidney organoids face challenges in achieving expression of LOH-specific genes.^45^ Hence, there is utility in using animal models to assemble a working genetic framework that could enrich for LOH development in culture conditions.

A compelling TFAP2A GRN candidate that warrants further investigation is KCTD1, which shares greater than 78% sequence homology with KCTD15.^46^ KCTD1 can directly inhibit TFAP2A as well as suppress canonical Wnt signaling, exhibiting congruence with known KCTD15 molecular mechanisms.^47, 48^ Human KCTD1 mutations cause scalp-ear-nipple (SEN) syndrome, which involves the formation of toxic amyloid-like aggregates thought to influence disease state.^49^ With the advent of prenatal screening and genotyping strategies, KCTD1 lesions have been linked to renal hypoplasia.^50^ Along these lines, mice harboring a Kctd1 mutation die perinatally due to kidney failure associated with defective ion homeostasis. Transcriptional profiling of these mutant kidneys identified 102 significantly upregulated genes, including known TFAP2A targets.^51^ TFAP2A and TFAP2B null mice also manifest lethal kidney defects.^52, 53^ Ultimately, our genetic studies elucidating TFAP2-KCTD network mechanisms in developing zebrafish nephrons can catalyze candidate identification and refine prenatal screening for renal defects.

## Author contributions

B.E.C. and R.A.W. designed the experiments. B.E.C., E.G.C., and A.G. conducted the experiments.

B.E.C., E.G.C., A.G., and R.A.W. analyzed the experiments. B.E.C. and R.A.W. wrote the manuscript.

## Supplemental Materials

**Fig. S1.**
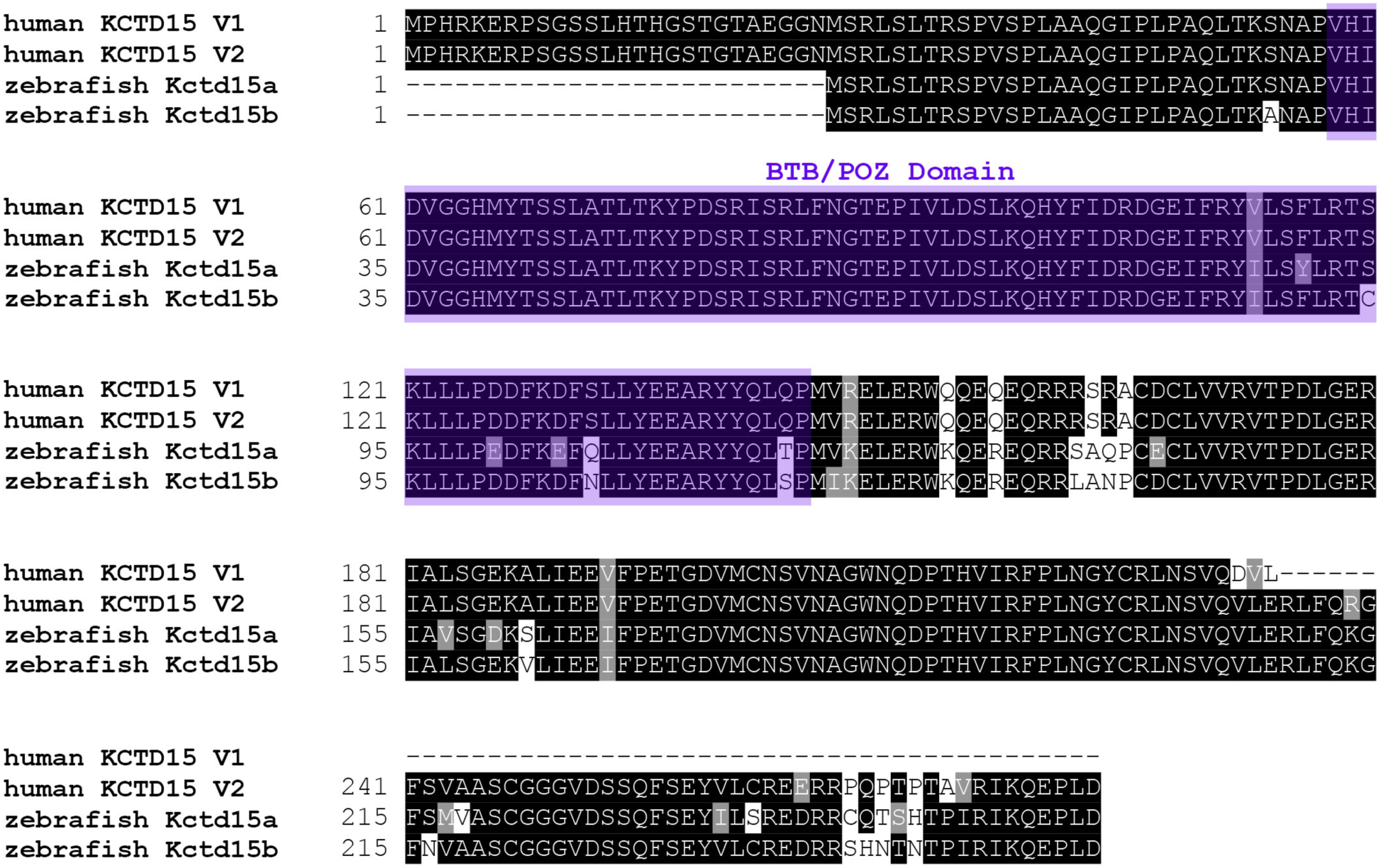
Zebrafish Kctd15a and Kctd15b paralogs exhibit high amino acid sequence conservation with human KCTD15.

**Fig. S2.**
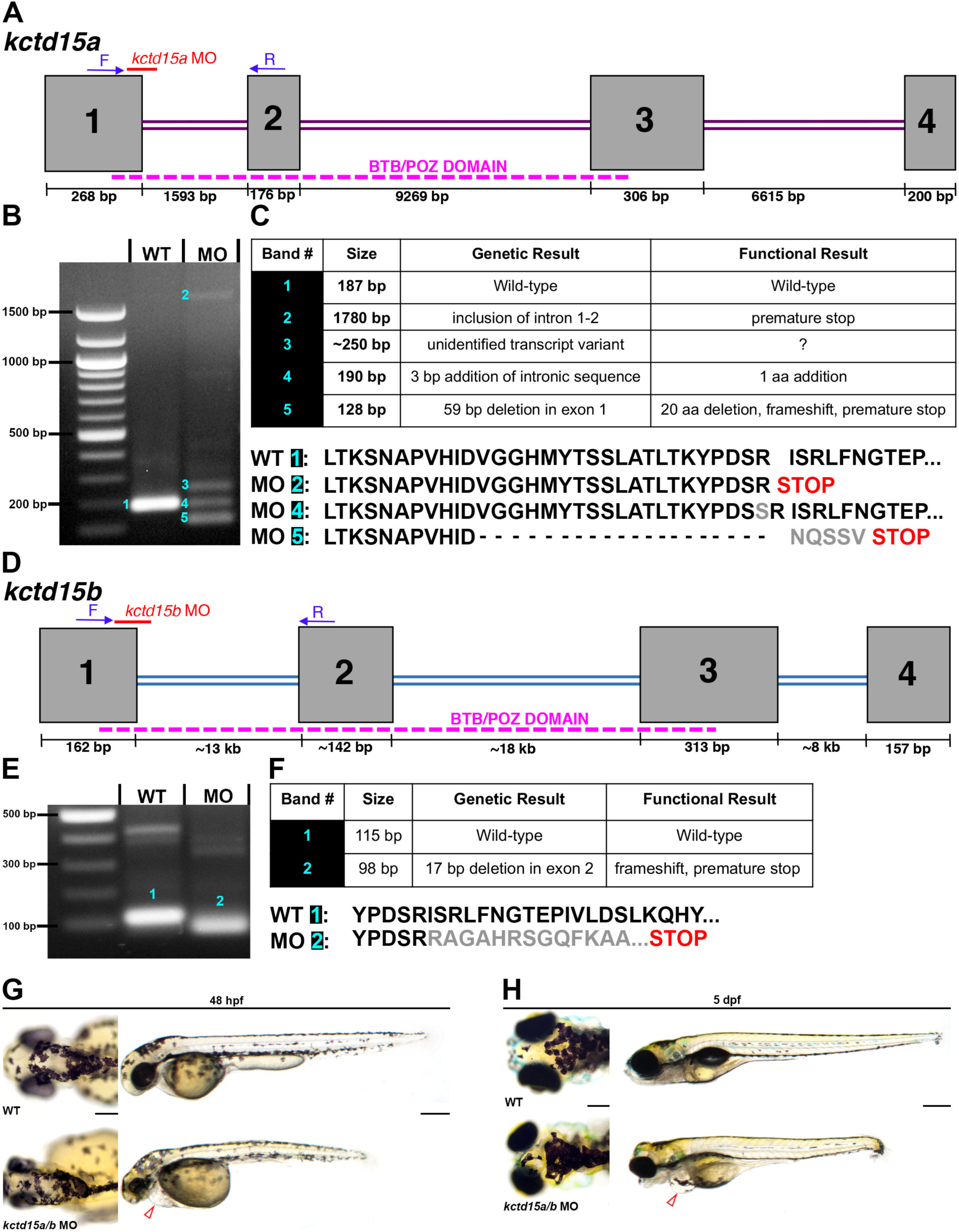
RT-PCR analysis confirms *kctd15a* and *kctd15b* MO reagents effectively block splicing.

**Fig. S3.**
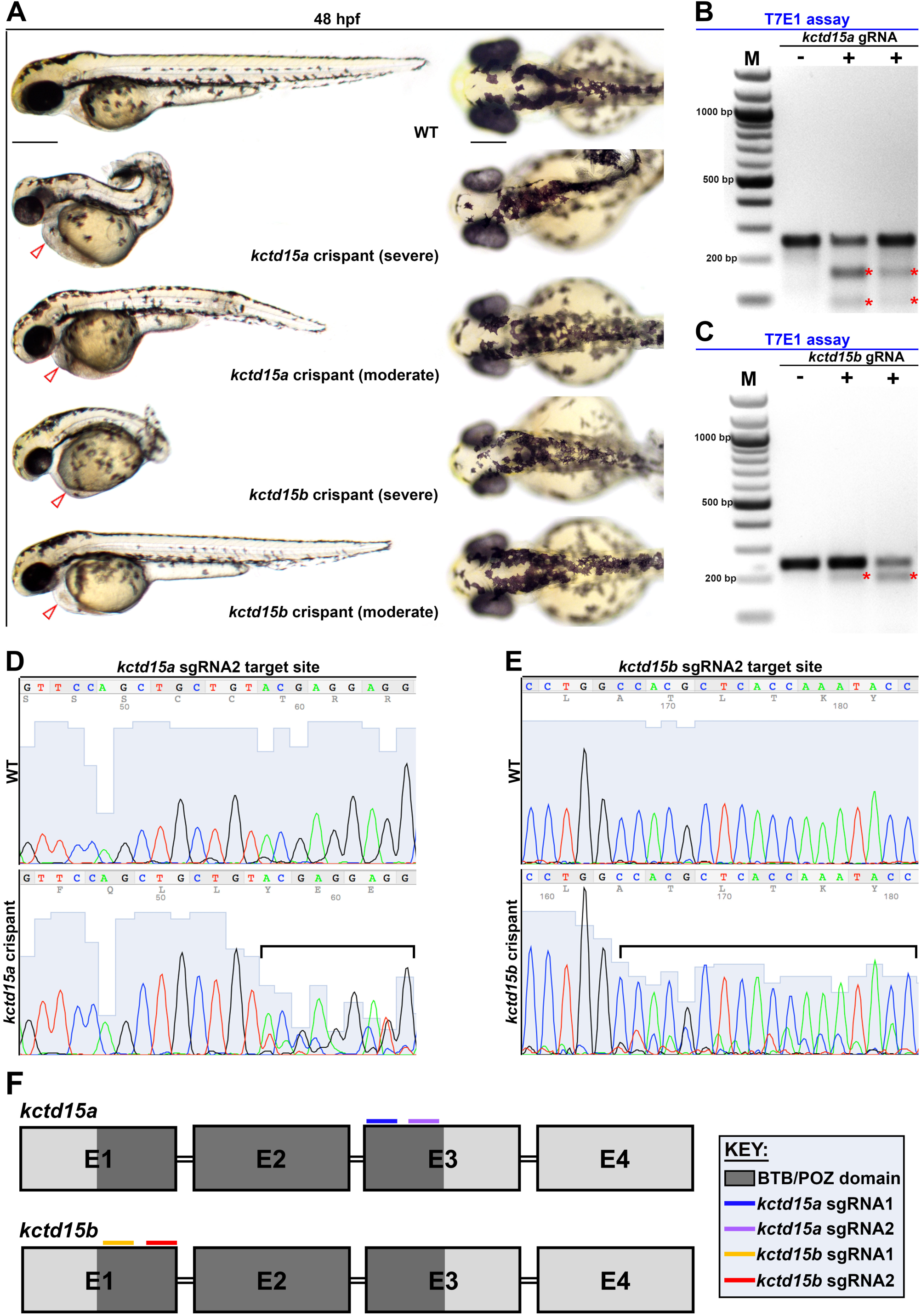
k*c*td15a and *kctd15b* gRNAs edit the genome verifying successful crispant generation.

**Fig. S4.**
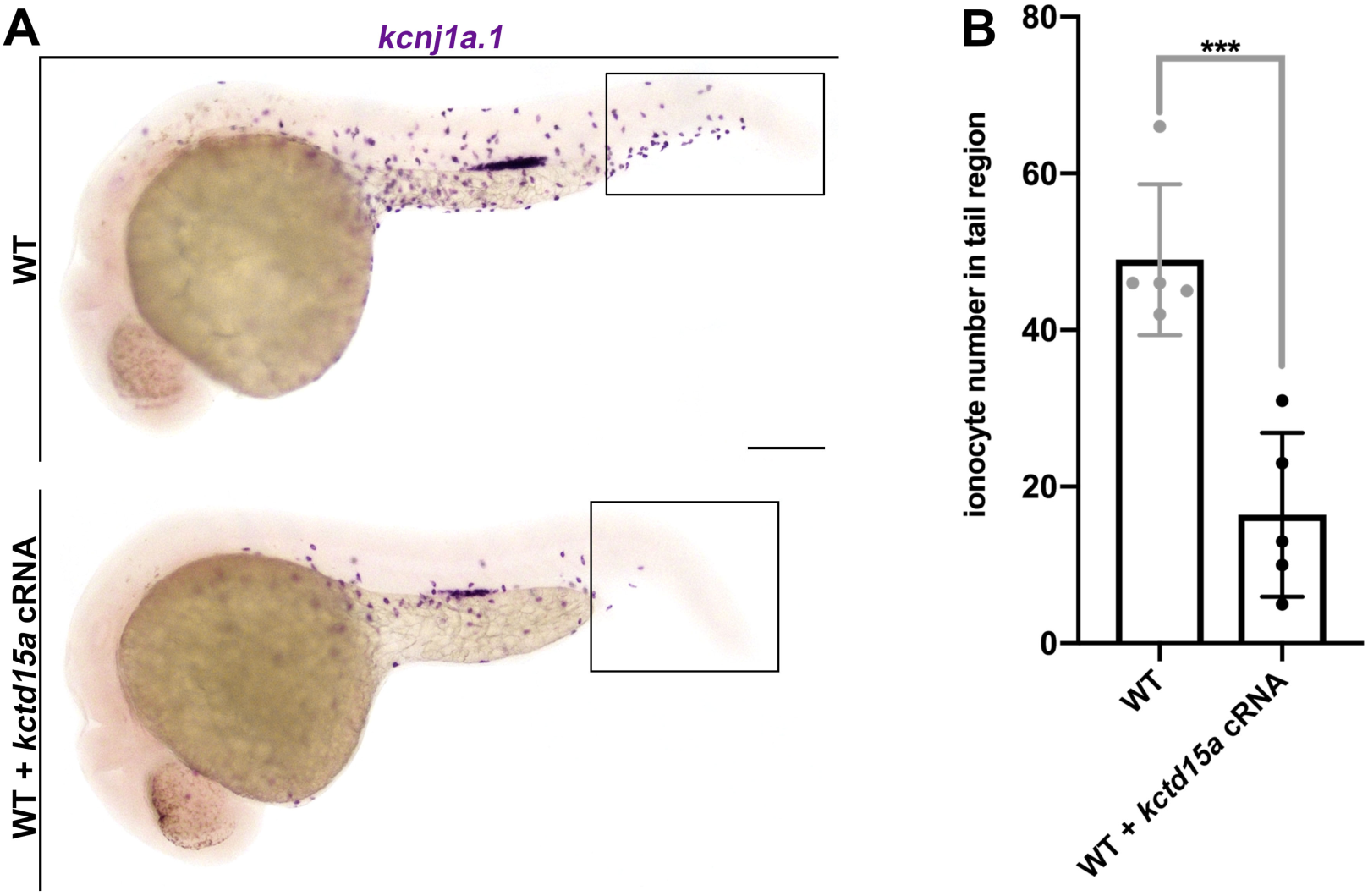
k*c*td15a overexpression represses ionocyte development.

**Table S1.**
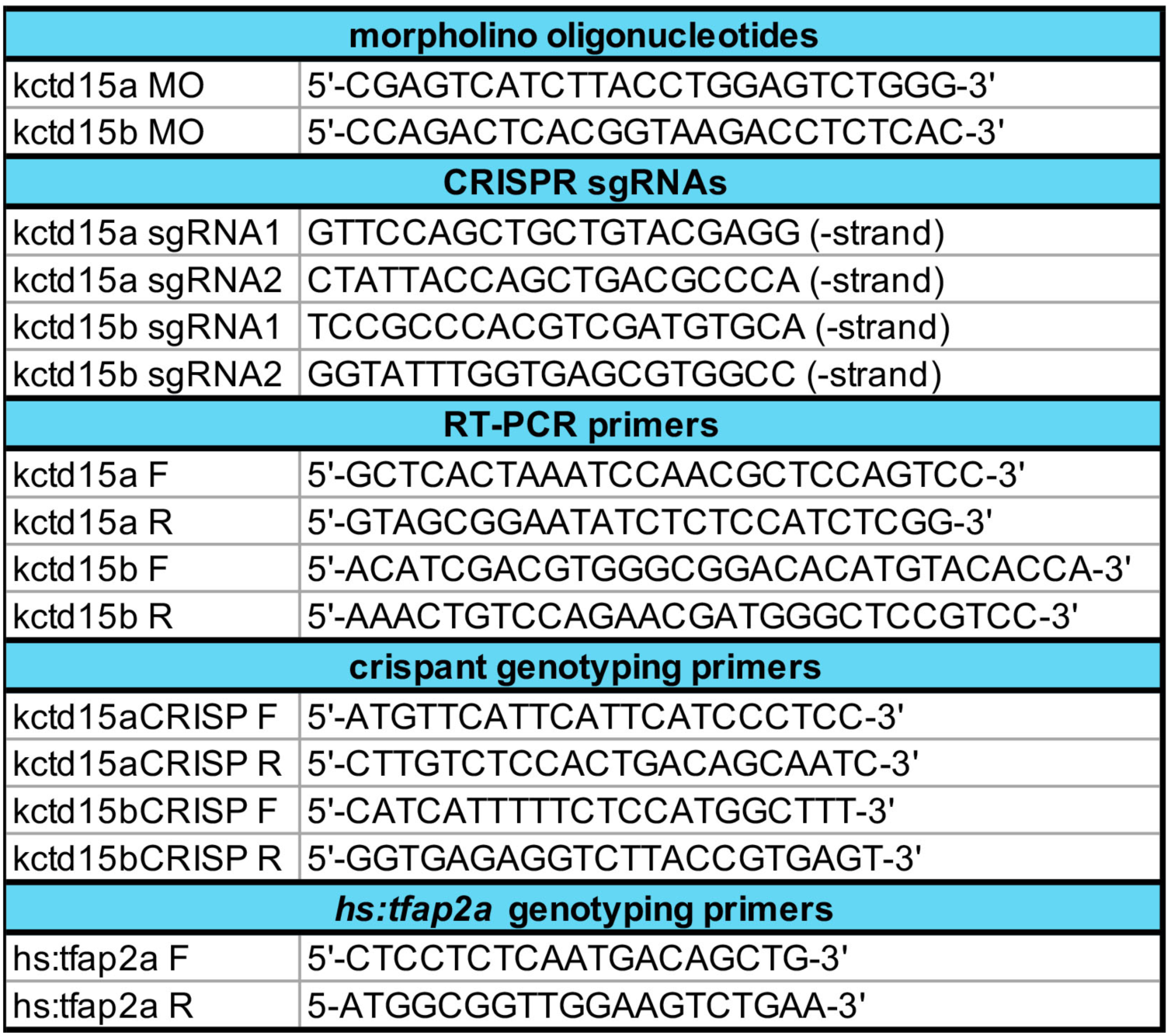
Compilation of primer, morpholino oligonucleotide, and CRISPR sgRNA sequences.

